# Perceived predation risk affects the development of among-individual behavioral variation in a naturally clonal freshwater fish

**DOI:** 10.1101/2023.11.25.568653

**Authors:** U Scherer, K Laskowski, M Kressler, S Ehlman, M Wolf, D Bierbach

## Abstract

Predation risk is a key driver of natural selection, influencing various aspects of prey behavior. While many studies focus on how predation risk affects average behavior at population level, less attention has been given to its potential impact on behavioral variation within prey populations. Here, we investigate the effect of perceived predation risk on among-individual behavioral variation in naturally clonal Amazon mollies. Juveniles were raised in two groups: one exposed to a predator during feeding (visual cues only) and the other one serving as a control group. We observed activity and feeding behavior (time spent feeding, visits to feeding spot) over a four-week period. (I) Individuals in the predator-exposed group were on average less active but there was no difference in average feeding behavior between the two groups, suggesting individuals strategically respond to threats based on behavior-specific cost-benefit trade-offs. (II) Among-individual behavioral variation was affected by perceived predation risk: in the absence of the predator, individuals developed pronounced differences in the time spent feeding while no such development was observed in the predator-exposed group. This result has the potential of affecting a wide range of fitness-relevant intraspecific interactions if lower among-individual feeding variation translate into reduced sizes differences. The presence of the predator initially reduced among-individual variation in activity and visits to the feeding spot, but these differences did not persist over time. Our findings highlight the importance of considering both population-level and individual-level responses to predation risk for a more comprehensive understanding of its ecological and evolutionary consequences.

## INTRODUCTION

Predation risk is one of the major forces of natural selection shaping various aspects of prey phenotypes including behavioral, morphological, and life-history traits (Chivers et al., 2008; Lima & Dill, 1990; Reznick & Endler, 1982; Walls et al., 1990). At the level of behavior, a labile trait that contains both relatively fixed (e.g., stereotypic) as well as more plastic components (Kelley & Magurran, 2003), predation risk has profound consequences, for instance, on grouping, movement, mating, and foraging (Bierbach et al., 2011; Doran et al., 2022; Sih, 1994; Sosna et al., 2019; Walls et al., 1990). To give two concrete examples, poecilid fish from high predation sides are bolder, i.e., they emerged faster from a shelter to explore a novel environment, than their conspecifics from low-predation sides (Brown et al., 2007; Harris et al., 2010) and, with increasing predation risk, reef fish (several species, including, e.g., parrotfish, *Sparisoma aurofrenatum* and surgeonfish, *Acanthurus bahianus*) consume drastically less food (approx. 90%) but fed at a faster rate (approx. 26%) (Catano et al., 2016).

The above examples illustrate that, up to now, the vast majority of studies have focused on how predation risk influences average behavior at population level. In contrast, and despite the ecological and evolutionary significance of among-individual behavioral differences (Bell et al., 2009; Bolnick et al., 2011; Carere & Gherardi, 2013; Ingley & Johnson, 2014; Réale et al., 2007; Wolf & Weissing, 2012), the effects predators might have on variation expressed among individuals within prey populations are hugely understudied (but see Arlinghaus et al., 2017; Dingemanse et al., 2009; Sommer-Trembo et al., 2016). The few existing studies addressing how predation risk affects among-individual behavioral variation provide inconclusive results. For example, wild-caught (i.e., predator-experienced) guppies, *Poecilia reticulata*, have been shown to homogenize regarding their risk-taking behavior when presented with a computer-animated predator, in contrast, the same study found that laboratory-bred (i.e., predator-naïve) guppies showed a diversification in behavior under the same testing conditions (Sommer-Trembo et al., 2016).

The discrepancy in results, i.e., diversification vs. homogenization of behavior in response to predation risk, may be routed in populations having diverse evolutionary histories (predator absence vs. predator presence, spatial-temporal fluctuations in predation pressure) leading to differences in genetic backgrounds (heterogenic populations where different genotypes may or may not react differently to predation risk vs. homogenic populations) (Barbosa et al., 2018; Dingemanse et al., 2009; Polverino et al., 2018). Furthermore, diversity in predator experience may play a vital role in changing an individual’s risk-perception and thus its reaction towards future predator encounters (Kelley & Magurran, 2003; Tulley & Huntingford, 1987). Additionally, in the wild, prey might be selected through lethal predator encounters that erase specific behavioral types from the population making it difficult to disentangle consumptive from non-consumptive effects (i.e., do observed patterns in behavioral variation stem from pre-selection though predators consuming certain individuals or do surviving individuals change their behavior according to their experience with predators?) (Arlinghaus et al., 2017). Therefore, to gain a better understanding of how predation risk influences behavioral variation within prey populations, we are in urgent need of studies that carefully control for evolutionary and experiential backgrounds as well as for other factors accecting variation like age, sex, size, and physiological state (Catano et al., 2016; Harris et al., 2010; Magnhagen & Borcherding, 2008).

In the current study, we thus used a highly-controlled experimental set-up to test the effect of perceived predation risk on the development of among-individual behavioral variation in natural clonal Amazon mollies, *Poecilia formosa*, seperated on day 15 of their life into otherwise near-identical environmental conditions: for the following 4 weeks, individuals were either reared in a predator treatment, where fish were forced to forage under simulated predation risk (i.e., a convict cichlid was presented during the feedings, visual contact only) or in a control treatment with no predator being present during feeding. In both treatments, we recorded individual activity (average swimming speed) and feeding behavior (time spent feeding and number of visits to the feeding spot) during feeding; two behaviors Amazon mollies are known to show strong individuality in (Bierbach et al., 2017; Laskowski et al., 2022; Scherer et al., 2023).

We predicted (I) the predator exposure to influence average behavior at population level, with fish in the predator treatment avoiding the predator and therefore being less active, spending less time feeding, and visiting the feeding spot less frequently compared to the control group. Furthermore, we predicted that (II) behavioral individuality in the three observed behaviors (activity, time spent feeding, visits to the feeding spot) would be affected by the predator exposure. However, since both diversification and homogenization of prey fish behavior in response to perceived predation risk are plausible, we did not make a directional prediction here. More specifically, one might expect that perceived predation risk leads to a diversification in behavior if individuals respond differently to potential risk (Huntingford & Wright, 1993; Magurran, 1990). Alternatively, perceived predation risk might homogenize behavior if individuals agree in their perception of risk and converge on an optimal behavioral response (Sommer-Trembo et al., 2016).

## METHODS

### Study species and animal care

The Amazon molly is a gynogenetically reproducing freshwater fish from the subtropics of North America (Lamatsch et al., 2005; Lampert & Schartl, 2008; Schlupp, 2005; Stöck et al., 2010; Warren et al., 2018) and the first discovered clonal vertebrate (Hubbs & Hubbs, 1932; Schultz, 1973), stemming from a single hybridization event between a female Atlantic molly (*Poecilia mexicana*) and male sailfin molly (*Poecilia latipinna*) about 100,000 years ago. Gynogenesic reproduction means that Amazon mollies require sperm from one of their parental species to trigger embryonic development, but paternal genetic material is not incorporated into the egg (Lampert & Schartl, 2008; Stöck et al., 2010; Warren et al., 2018) except of rare cases of male DNA fragment introgression (Kallman, 1962; Rasch et al., 1965; Turner et al., 1980). Resulting offspring are therefore genetically identical to their mother and each other.

Our experimental fish were lab-reared descendants of wild-caught fish originating from waters around the Mexican city of Tampico. Regular molecular checks confirmed that all *P. formosa* individuals are clones. Breeding of *P. formosa* was done with male *P. mexicana* individuals in large mixed-age uni-clonal tanks. All fish were fed twice a day with flake food (TetraMin) and once a week with live or frozen *Chironomid* larvae. Temperature was maintained at 25°C with a 12/12 light/dark circle provided.

### Experimental procedure

We isolated several gravid females into smaller tanks equipped with artificial plants and the same water conditions and feeding regime as outlined above. After they gave birth, we immediately removed the mothers and raised offspring in sibling groups for 2 weeks on *Artemia nauplii* and dusted flakefood (TetraMin). We then placed offspring from *N* = 6 mothers individually in each of 48 experimental tanks. After 2 days of habituation during which the feeding of the first two weeks was continued, we started our experimental feeding trials. Half of the individuals (*N* = 24) were presented with a water-filled bottle at which we clamped a food tablet (‘empty’ control treatment). The other half (*N* = 23) was presented with the same bottle and food tablet but this time, the bottle held a convict cichlid (*Amatitlania nigrofasciata,* predator treatment). For each trial, predators were chosen randomly from a pool of 100 juvenile cichlids (200 L tank, holding conditions as above). Bottles were put into the tanks every Monday, Wednesday, and Friday for 4 weeks so that every fish was presented 12 times with either an empty or a predator bottle. After the bottle with the food tablet was inserted, we gave our fish 30 min to feed and recorded at 2 frames per second from above (Laskowski et al., 2022). On days without feeding trials, fish were fed in the morning with a food tablet and the remaining food was removed in the evening. Feeding trials were tracked using software Ethovision XT (Noldus Inc.). We scored how many times the fish visited the food tablet (‘number of visits to feeding spot’), the time spend feeding on the tablet (‘feeding duration’, in %) as well as average velocity (‘activity’, in cm/sec).

### Statistical analyses

#### General details

Statistical analyses were performed in R 4.2.1 (R Core Team, 2022). LMMs (linear mixed-effect models) were fitted using the *lmer*-package (Bates et al., 2015). Variance components (repeatabilities, among-, and within-individual variation) with 95% confidence intervals (CIs) were estimated from LMMs following (Hertel et al., 2020). We tested whether variance components differ significantly between treatments by comparing the 95% CIs (significant difference between treatment when CIs do not overlap). Significance for behavioral repeatabilities per se was derived from the 95% CIs being distinctly different to zero. Repeatability is the most widely used statistic to test for behavioral individuality and to quantify how much of the total variation observed is due to differences among individuals (Bell et al., 2009). Model assumptions were verified using q-q plots and residual plots. Most parsimonious models were selected via stepwise backward removal of insignificant terms.

Our three target variables (activity, feeding duration, visits to feeding spot) were moderately (activity and feeding duration), weakly (activity and visits to feeding spot), or not (feeding duration and visits to feeding spot) correlated with each other (for more information and R^2^ see **Supplementary Table S1**), we therefore did not consider these variables redundant.

#### (I) Does the predator exposure affect population level average behavior?

We tested if average behavior differs between the predator and control treatment by building LMMs with the behavior of interest as response (activity, feeding duration, and visits to feeding spot, respectively) and treatment (predator vs. control) as fixed effect. We further included trial number (1-12) as fixed effect to account for potential effects of time (e.g., fish getting older or habituating over the course of the experiment), as well as the trial number-treatment interaction term to account for potential differences between the two treatments in how behavior changes over time. We included random intercepts for both individuals and families (*N* (predator and control) = 559 observations from 47 individuals per model).

#### (II) Does the predator exposure affect behavioral variation?

To test if perceived predation risk affects behavioral individuality, we estimated repeatability, among- and within-individual variation for each of our three target behaviors (activity, feeding duration, feeding bouts) in the two treatments (predator vs. control) separately. That is, for each behavior and treatment, we ran an LMM with the behavior of interest as response and experimental week (1-4) as fixed effect (in total 6 models, for each behavior: *N* (control) = 285 observations from 24 individuals and *N* (predator) = 274 observations from 23 individuals). As random terms, we included individual intercepts and individual slopes as well intercepts for family. In order to assess how variance components develop over the course of the experiment, we ran each of the above 6 models 4 times, during each run, we centered the model to a different week. That is, we ‘sliced’ the model at different time points allowing us to pull variance estimates for each week (Laskowski et al., 2022).

## RESULTS

### (I) Fish were less active in the presence of the predator, but no effect on feeding behavior

Fish in the predator treatment were generally less active than fish in the control treatment (estimate [CI] = -0.390 [-0.751--0.028], *p* = 0.035; **Supplementary Table S2, Figure 1a**;). And in both treatments, individuals decreased their activity over the course of the experiment (estimate [CI] = -0.147 [-0.170--0.125], *p* <0.001; **Supplementary Table S2,** **Figure 1a**). There was no difference in feeding duration (**Figure 1b**) and in the number of visits to the feeding spot (**Figure 1c**) between the predator and control treatment (**Supplementary Table S2**), but both behaviors changed over the course of the experiment: fish increased their time spent feeding (estimate [CI] = 0.022 [0.018-0.026], *p* <0.001; **Supplementary Table S2,** **Figure 1b**), and visited the feeding spot more often (estimate [CI] = 0.858 [0.369 – 1.347], *p* = 0.001; **Supplementary Table S2,** **Figure 1c**) when they got older.

**Figure 1.**
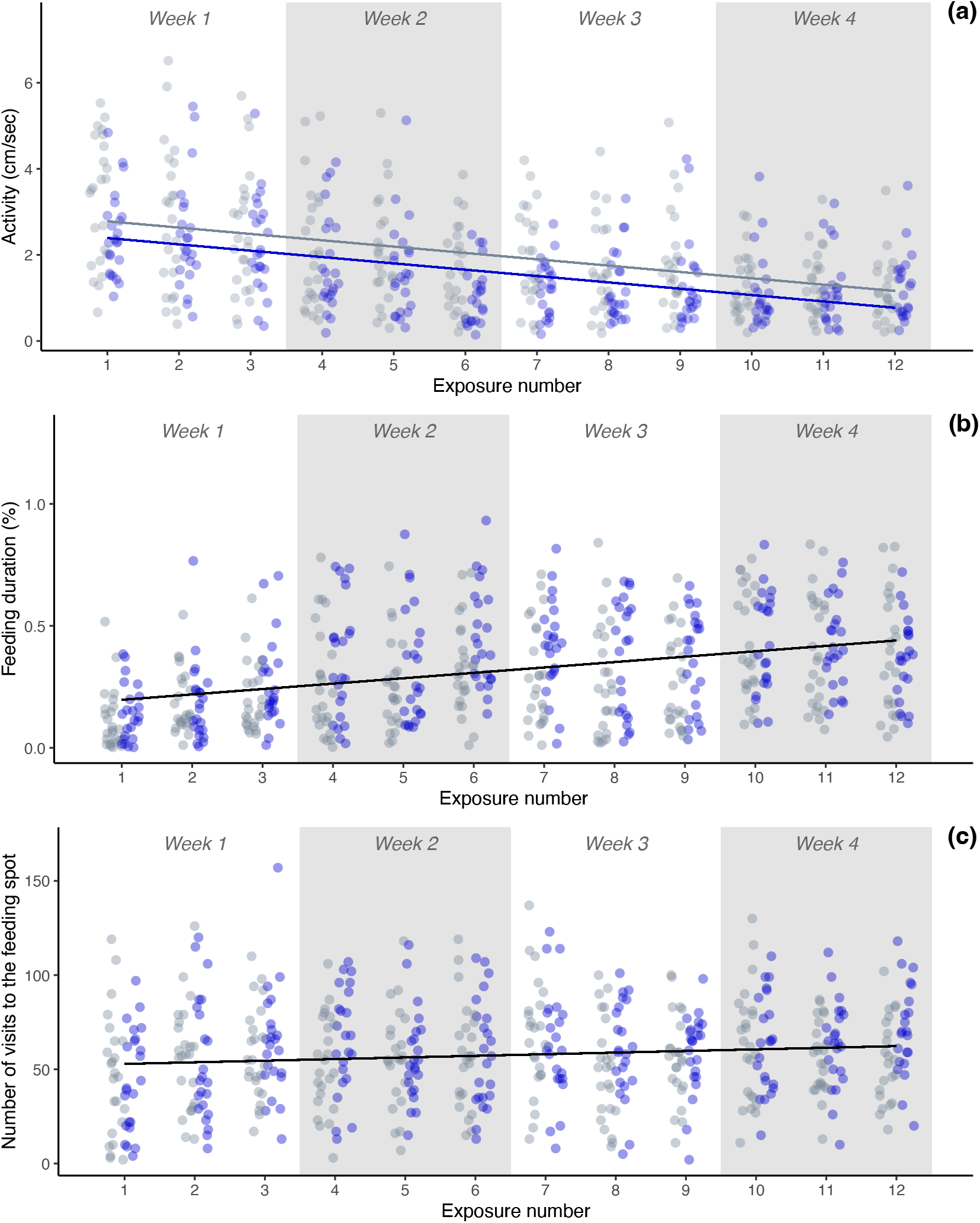
Effects of predator exposure on average behavior at population level over time: (a) In the presence of the predator (blue points), individuals were on average less active than in the control (gray points) and in both treatments, individuals became less active over time. (b-c) In both treatments, individuals increased their feeding duration and the number of visits to the feeding spot over time.

### (II) Predator exposure affected variation among but not within individuals

Over the course of the experiment, among-individual variation, and hence repeatability, in the time spent feeding increased significantly in the absence of the predator (i.e., in the control treatment). In the predator treatment, however, both variance components remained stable and significantly lower than in the control treatment (**Figure 2e-f**, **Supplementary Table S4**). The diversification in feeding in the control treatment is characterized by a fanning-out pattern of individual BLUPs (best linear unbiased predictors, i.e., random intercepts for individuals) over the four experimental weeks (**Figure 2d**). In the control treatment, we additionally observed higher initial among-individual variation (but not repeatability) in activity, compared to the predator treatment, which decreased significantly over the course of the experiment, resulting in similar patterns in both treatments towards the end of the experiment (**Figure 2a-b**, **Supplementary Table S3**). The homogenization of activity in the control treatment is characterized by a narrowing pattern of individual BLUPs over the four experimental weeks (**Figure 2d, h**). Regarding the number of visits to the feeding spot, we found among-individual variation and repeatability to be initially higher in the control compared to the predator treatment; but the effect was apparent in experimental week 1 and 2 only (**Figure 2i-j**, **Supplementary Table S5**). For all three behaviors (activity, feeding duration, visits to feeding spot), we found no differences in within-individual variation between the two treatments (**Figure 2c, g, k**, **Supplementary Tables S2-S5**).

**Figure 2.**
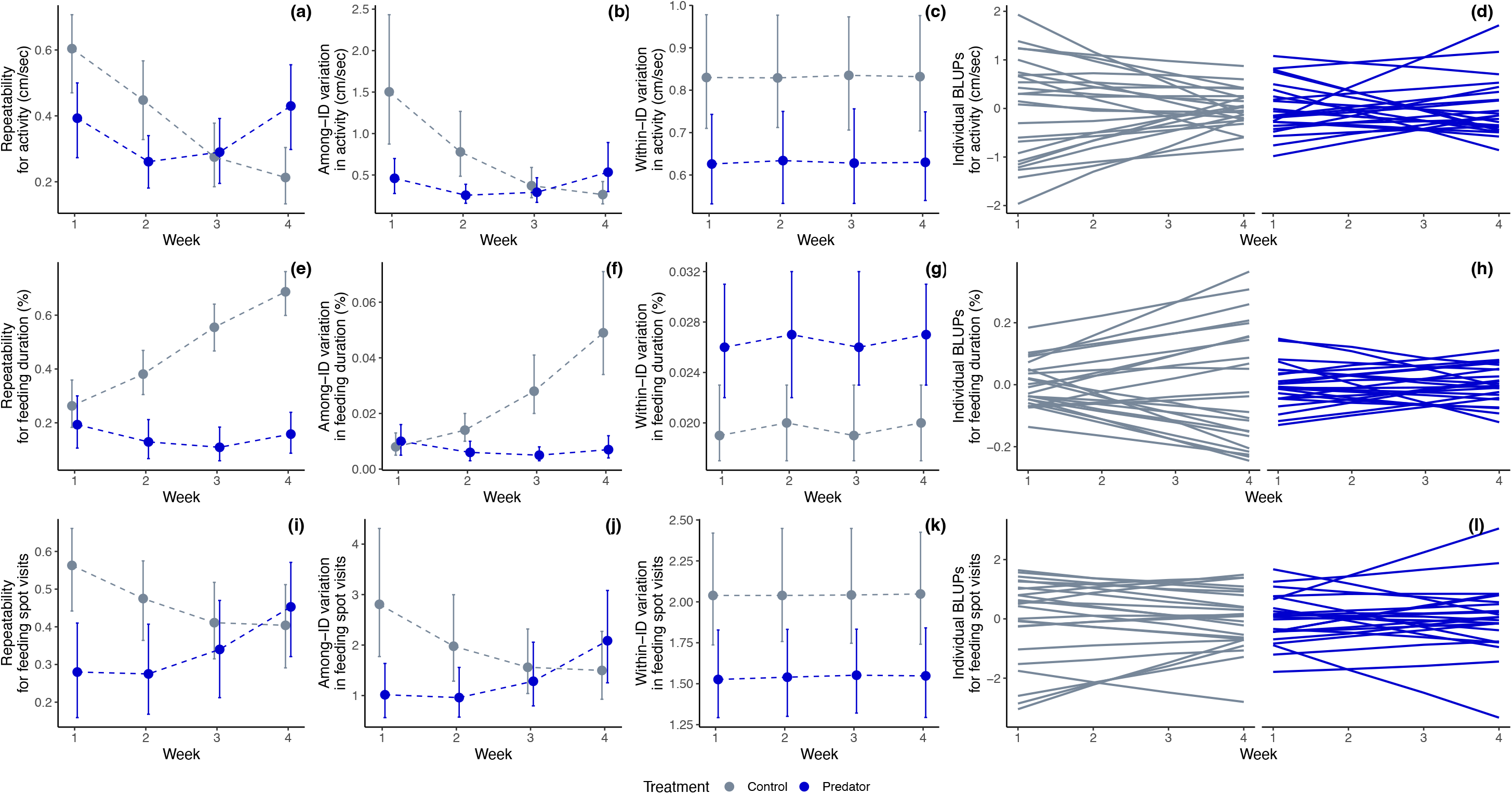
Variance components for individuals in the predator vs. control treatment, shown are (a, e, i) repeatabilities, (b, f, j) among-individual variation, (c, g, k) within-individual variation, and (d, h, l) individual BLUPs (best linear unbiased predictors, i.e., random intercepts for individuals).

## DISCUSSION

In summary, we found partial support for our first hypothesis (i.e., on average, individuals develop avoidance behavior towards the predator): fish were on average less active in the presence of the predator but there was no difference in average feeding behavior (time spent feeding and number of visits to the feeding spot). Regarding our second hypothesis (i.e., predation risk affects among-individual behavioral variation), we observed a significant diversification in the time spent feeding in the absence of the predator, i.e., individuals developed more pronounced among-individual variation, compared to fish that foraged under simulated predation risk. The presence of the predator inhibited among-individual variation in activity and in the visits to the feeding spot at first, but there were no differences at the end of the experiment. For activity, this pattern was associated with a homogenization of behavior, i.e., individuals become more similar to one another in the absence of the predator.

Individuals were on average less active in the presence of the predator but there was no difference in average feeding behavior (feeding duration and number of visits to the feeding spot). The predator’s lack of influence on feeding behavior is in line with the predation risk allocation hypothesis, which postulates that as exposure to predation risk increases, individuals reduce their avoidance of predators because the associated loss of energy intake becomes too significant (Lima & Bednekoff, 1999). Our fish showing predator avoidance by reducing their activity may indicate that the two behaviors differ in their cost-benefit trade-offs. More specifically, decreasing both activity and feeding can help minimize overall exposure to danger (Milinski, 1986; O’Connor et al., 2015; Scherer et al., 2017), however, feeding is crucial for energy intake and cannot be entirely avoided, whereas reducing activity is a less costly strategy.

While average time spent feeding did not differ among fish in the presence vs. absence of the predator, we observed strong among-individual diversification when the predator was absent and no substantial changes of among-individual variation in the presence of the predator over time. A potential explanation of the observed pattern could be that our two treatments differ in environmental uncertainty. That is, in the predator treatment, the sheer presence of the predator might be a very accurate cue regarding potential threat leading to high agreement in the assessment of risk and leading to individuals conforming to an ‘optimal response’ (Cooper & Blumstein, 2015; Sommer-Trembo et al., 2016). In the absence of the predator, on the other hand, individuals must rely on more inaccurate cues that might lead to more pronounced differences in how individuals perceive and assess the risk. For example, the performance of feeding trials is necessarily associated with a disturbance (the feeding apparatus is put in and taken out of the tank), which may or may not be perceived as a potential threat. Likewise, the potential danger associated with the feeding apparatus itself might be interpreted differently by different individuals. Future studies may investigate the effect of environmental uncertainty and cue reliability on among-individual behavioral variation. It will be particularly interesting, to focus on feedback loops as a mechanism underlying the development of among-individual differences (Ehlman et al., 2022).

Fish that foraged under simulated predation risk initially showed lower among-individual variation in activity and in visits to the feeding spot compared to the predator free-scenario, but these differences did not persist. Similar to our proposed explanation for lower among-individual variation in how much time fish spend feeding in the presence of the predator, this pattern could be caused by individuals aiming to exhibit an optimal response when facing a potential predatory threat (see above). The effect (i.e., lower among-individual variation in the presence compared to the absence of a predator) may appear only initially due to other developmental processes, e.g., for activity, we observed substantial homogenization of behavior in the predator-free scenario.

The inhibition of the development of among-individual diversity in feeding behavior as a response to perceived predation risk could have substantial ecological and evolutionary consequences. Specifically, decreased among-individual variation in feeding behavior might translate into differences in among-individual growth and body size, which in turn could have implications on a wide range of fitness-relevant intraspecific interactions, including competition and hierarchies (Buston & Cant, 2006; Smith & Brown, 1986), and reproductive behaviors (Bisazza & Marconato, 1988), as well as interspecific interactions, e.g., by affecting prey oddity and consequently predator hunting success (Landeau & Terborgh, 1986). While we can here only speculate about the consequences of predator induced homogenization of prey behavior, future studies may investigate this in more detail.

In conclusion, our results support the risk allocation hypothesis and indicate that individuals make strategic decisions about how to avoid a potential threat depending on the specific trade-offs (i.e., no predator avoidance when avoidance is too costly). Moreover, perceived predation risk suppressed among-individual diversity, particularly evident in the time spent feeding, with initial impacts on activity and visits to the feeding spot but without enduring effects. This suggests that when confronted with a predator, individuals align their risk evaluation, whereas in the absence of an immediate threat, variation in environmental perception among individuals become more pronounced. Overall, we show that, next to the effects predators have on average population level behavior, there are also important – and up to now largely overlooked – effects on variation among individuals, with potentially important ecological and evolutionary consequences.

