## Supplementary information for "Perceived predation risk affects the development of among-individual behavioral variation in a naturally clonal freshwater fish"

**Table S1** Linear mixed-effects models testing if our three target behaviors (activity, feeding duration, visits to feeding spot) were correlated: activity and feeding duration as well as activity and visits to the feeding spot were positively correlated but feeding duration and number of visits to the feeding spot were not correlated; tested for the predator (left models) and control (right models) treatment separately.

| Response | Predictors | Predator | | | Control | | |
|---|---|---|---|---|---|---|---|
| | | Estimates | CI | p | Estimates | CI | p |
| Activity (cm/sec) | (Intercept) | 2.779 | 2.541 – 3.017 | - | 3.208 | 2.887 – 3.530 | - |
| | Feeding duration (%) | -3.508 | -3.970 – -3.046 | <0.001 | -4.301 | -4.863 – -3.739 | <0.001 |
| | Random Effects | | | | | | |
| | $\sigma^2$ | 0.52 | | | 0.70 | | |
| | $\tau_{00}$ | 0.15 Fish ID | | | 0.42 Fish ID | | |
| | | 0.00 Family | | | 0.00 Family | | |
| | N | 23 Fish ID | | | 24 Fish ID | | |
| | | 6 Family | | | 6 Family | | |
| | Observations | 274 | | | 285 | | |
| | Marginal $R^2$ / Conditional $R^2$ | 0.520 / NA | | | 0.545 / NA | | |
| Activity (cm/sec) | (Intercept) | 0.982 | 0.581 – 1.383 | - | 1.133 | 0.738 – 1.528 | - |
| | Feeding spot visits | 0.010 | 0.005 – 0.015 | 0.001 | 0.015 | 0.009 – 0.021 | <0.001 |
| | Random Effects | | | | | | |
| | $\sigma^2$ | 0.94 | | | 1.25 | | |
| | $\tau_{00}$ | 0.10 Fish ID | | | 0.21 Fish ID | | |
| | | 0.04 Family | | | 0.00 Family | | |
| | ICC | 0.13 | | | | | |
| | N | 23 Fish ID | | | 24 Fish ID | | |
| | | 6 Family | | | 6 Family | | |
| | Observations | 274 | | | 285 | | |
| | Marginal $R^2$ / Conditional $R^2$ | 0.063 / 0.189 | | | 0.113 / NA | | |
| Feeding duration (%) | (Intercept) | 0.329 | 0.224 – 0.434 | - | 0.269 | 0.183 – 0.355 | - |
| | Feeding spot visits | 0.001 | -0.000 – 0.002 | 0.281 | 0.000 | -0.001 – 0.001 | 0.543 |
| | Random Effects | | | | | | |
| | $\sigma^2$ | 0.03 | | | 0.03 | | |
| | $\tau_{00}$ | 0.00 Fish ID | | | 0.01 Fish ID | | |
| | | 0.01 Family | | | 0.00 Family | | |
| | ICC | 0.28 | | | 0.33 | | |
| | N | 23 Fish ID | | | 24 Fish ID | | |
| | | 6 Family | | | 6 Family | | |
| | Observations | 274 | | | 285 | | |
| | Marginal $R^2$ / Conditional $R^2$ | 0.005 / 0.287 | | | 0.001 / 0.327 | | |

**Table S2** Linear mixed-effects models testing if individuals that were forced to forage in the presence of the predator behaved differently (regarding their activity, time spent feeding, and visits to the feeding spot) compared individuals in the control treatment. We considered a potential effect of the trial number and the possibility that the trial number may affect behavior differently in the two treatments. Shown are full models (left) containing all predictors and final models (right) containing significant predictors only.

| | | Full model | | | Final model | | |
|---|---|---|---|---|---|---|---|
| **Response** | **Predictors** | **Estimates** | **CI** | **p** | **Estimates** | **CI** | **p** |
| Activity (cm/sec) | (Intercept) | 3.043 | 2.719 – 3.366 | - | 2.929 | 2.638 – 3.220 | - |
| | Treatment [Predator] | -0.622 | -1.084 – -0.160 | - | -0.390 | -0.751 – -0.028 | 0.035 |
| | Trial no. | -0.165 | -0.196 – -0.134 | - | -0.147 | -0.170 – -0.125 | <0.001 |
| | Treatment × Trial no. | 0.036 | -0.009 – 0.080 | 0.114 | - | - | - |
| | **Random Effects** | | | | | | |
| | $\sigma^2$ | 0.85 | | | 0.85 | | |
| | $\tau_{00}$ | 0.33 Fish ID | | | 0.33 Fish ID | | |
| | | 0.00 Family | | | 0.00 Family | | |
| | N | 47 Fish ID | | | 47 Fish ID | | |
| | | 6 Family | | | 6 Family | | |
| | Observations | 559 | | | 559 | | |
| | Marginal $R^2$ / Conditional $R^2$ | 0.260 / NA | | | 0.257 / NA | | |
| Feeding duration (%) | (Intercept) | 0.143 | 0.063 – 0.223 | - | 0.175 | 0.104 – 0.246 | - |
| | Treatment [Predator] | 0.065 | -0.014 – 0.145 | - | - | - | - |
| | Exposure no. | 0.023 | 0.018 – 0.029 | - | 0.022 | 0.018 – 0.026 | <0.001 |
| | Treatment × Exposure no. | -0.003 | -0.011 – 0.005 | 0.478 | - | - | - |
| | **Random Effects** | | | | | | |
| | $\sigma^2$ | 0.03 | | | 0.03 | | |
| | $\tau_{00}$ | 0.01 Fish ID | | | 0.01 Fish ID | | |
| | | 0.00 Family | | | 0.00 Family | | |
| | ICC | 0.35 | | | 0.36 | | |
| | N | 47 Fish ID | | | 47 Fish ID | | |
| | | 6 Family | | | 6 Family | | |
| | Observations | 559 | | | 559 | | |
| | Marginal $R^2$ / Conditional $R^2$ | 0.138 / 0.439 | | | 0.126 / 0.439 | | |
| Visits to feeding spot | (Intercept) | 52.676 | 43.337 – 62.016 | - | 52.001 | 44.675 – 59.326 | - |
| | Treatment [Predator] | -1.372 | -13.282 – 10.538 | - | - | - | - |
| | Exposure no. | 0.466 | -0.218 – 1.149 | - | 0.858 | 0.369 – 1.347 | 0.001 |
| | Treatment × Exposure no. | 0.799 | -0.177 – 1.774 | 0.109 | - | - | - |
| | **Random Effects** | | | | | | |
| | $\sigma^2$ | 407.54 | | | 409.62 | | |
| | $\tau_{00}$ | 273.62 Fish ID | | | 276.86 Fish ID | | |
| | | 22.19 Family | | | 22.34 Family | | |
| | ICC | 0.42 | | | 0.42 | | |
| | N | 47 Fish ID | | | 47 Fish ID | | |
| | | 6 Family | | | 6 Family | | |
| | Observations | 559 | | | 559 | | |
| | Marginal $R^2$ / Conditional $R^2$ | 0.020 / 0.432 | | | 0.012 / 0.429 | | |

**Table S3** Variance components for activity in the control vs. predator treatment.

| Behavior | Variance component | Week | Control | | | Predator | | |
|---|---|---|---|---|---|---|---|---|
| | | | Estimate | Lower CI | Upper CI | Estimate | Lower CI | Upper CI |
| Activity (cm/sec) | Repeatability for individual | 1 | 0.604 | 0.470 | 0.707 | 0.393 | 0.273 | 0.500 |
| | Repeatability for family | | 0.021 | 0.005 | 0.059 | 0.000 | 0.000 | 0.000 |
| | Among-individual variation in intercepts | | 1.503 | 0.873 | 2.433 | 0.460 | 0.277 | 0.699 |
| | Among-individual variation in slopes | | 0.103 | 0.076 | 0.132 | 0.084 | 0.055 | 0.121 |
| | Among-family variation in intercepts | | 0.052 | 0.012 | 0.143 | 0.000 | 0.000 | 0.000 |
| | Within-individual variation | | 0.830 | 0.710 | 0.978 | 0.626 | 0.532 | 0.743 |
| | Repeatability for individual | 2 | 0.448 | 0.328 | 0.567 | 0.261 | 0.181 | 0.340 |
| | Repeatability for family | | 0.010 | 0.002 | 0.029 | 0.000 | 0.000 | 0.000 |
| | Among-individual variation in intercepts | | 0.778 | 0.485 | 1.267 | 0.256 | 0.159 | 0.388 |
| | Among-individual variation in slopes | | 0.108 | 0.073 | 0.156 | 0.093 | 0.056 | 0.140 |
| | Among-family variation in intercepts | | 0.018 | 0.004 | 0.053 | 0.000 | 0.000 | 0.000 |
| | Within-individual variation | | 0.829 | 0.712 | 0.977 | 0.634 | 0.533 | 0.750 |
| | Repeatability for individual | 3 | 0.275 | 0.185 | 0.378 | 0.290 | 0.195 | 0.392 |
| | Repeatability for family | | 0.013 | 0.003 | 0.036 | 0.000 | 0.000 | 0.000 |
| | Among-individual variation in intercepts | | 0.371 | 0.226 | 0.591 | 0.292 | 0.170 | 0.467 |
| | Among-individual variation in slopes | | 0.126 | 0.077 | 0.193 | 0.089 | 0.056 | 0.138 |
| | Among-family variation in intercepts | | 0.017 | 0.005 | 0.051 | 0.000 | 0.000 | 0.000 |
| | Within-individual variation | | 0.835 | 0.706 | 0.973 | 0.628 | 0.533 | 0.756 |
| | Repeatability for individual | 4 | 0.213 | 0.133 | 0.304 | 0.430 | 0.298 | 0.555 |
| | Repeatability for family | | 0.013 | 0.003 | 0.032 | 0.000 | 0.000 | 0.000 |
| | Among-individual variation in intercepts | | 0.264 | 0.152 | 0.421 | 0.534 | 0.300 | 0.892 |
| | Among-individual variation in slopes | | 0.127 | 0.071 | 0.204 | 0.077 | 0.053 | 0.112 |
| | Among-family variation in intercepts | | 0.016 | 0.004 | 0.042 | 0.000 | 0.000 | 0.000 |
| | Within-individual variation | | 0.832 | 0.704 | 0.976 | 0.630 | 0.540 | 0.749 |

**Table S4** Variance components for time spent feeding in the control vs. predator treatment.

| Behavior | Variance component | Week | Control | | | Predator | | |
|---|---|---|---|---|---|---|---|---|
| | | | Estimate | Lower CI | Upper CI | Estimate | Lower CI | Upper CI |
| Feeding duration (%) | Repeatability for individual | 1 | 0.263 | 0.183 | 0.359 | 0.193 | 0.106 | 0.300 |
| | Repeatability for family | | 0.000 | 0.000 | 0.000 | 0.266 | 0.092 | 0.463 |
| | Among-individual variation in intercepts | | 0.008 | 0.005 | 0.013 | 0.010 | 0.005 | 0.016 |
| | Among-individual variation in slopes | | 0.003 | 0.002 | 0.005 | 0.001 | 0.001 | 0.002 |
| | Among-family variation in intercepts | | 0.000 | 0.000 | 0.000 | 0.014 | 0.004 | 0.032 |
| | Within-individual variation | | 0.019 | 0.017 | 0.023 | 0.026 | 0.022 | 0.031 |
| | Repeatability for individual | 2 | 0.381 | 0.305 | 0.469 | 0.129 | 0.067 | 0.212 |
| | Repeatability for family | | 0.000 | 0.000 | 0.000 | 0.255 | 0.098 | 0.454 |
| | Among-individual variation in intercepts | | 0.014 | 0.010 | 0.020 | 0.006 | 0.003 | 0.010 |
| | Among-individual variation in slopes | | 0.003 | 0.002 | 0.005 | 0.001 | 0.001 | 0.002 |
| | Among-family variation in intercepts | | 0.000 | 0.000 | 0.000 | 0.012 | 0.004 | 0.027 |
| | Within-individual variation | | 0.020 | 0.017 | 0.023 | 0.027 | 0.022 | 0.032 |
| | Repeatability for individual | 3 | 0.555 | 0.467 | 0.641 | 0.109 | 0.059 | 0.184 |
| | Repeatability for family | | 0.000 | 0.000 | 0.000 | 0.264 | 0.089 | 0.435 |
| | Among-individual variation in intercepts | | 0.028 | 0.020 | 0.041 | 0.005 | 0.003 | 0.008 |
| | Among-individual variation in slopes | | 0.003 | 0.002 | 0.004 | 0.001 | 0.001 | 0.002 |
| | Among-family variation in intercepts | | 0.000 | 0.000 | 0.000 | 0.012 | 0.003 | 0.026 |
| | Within-individual variation | | 0.019 | 0.017 | 0.023 | 0.026 | 0.023 | 0.032 |
| | Repeatability for individual | 4 | 0.687 | 0.599 | 0.762 | 0.158 | 0.087 | 0.239 |
| | Repeatability for family | | 0.000 | 0.000 | 0.000 | 0.246 | 0.096 | 0.409 |
| | Among-individual variation in intercepts | | 0.049 | 0.034 | 0.071 | 0.007 | 0.004 | 0.012 |
| | Among-individual variation in slopes | | 0.003 | 0.002 | 0.003 | 0.001 | 0.001 | 0.002 |
| | Among-family variation in intercepts | | 0.000 | 0.000 | 0.000 | 0.011 | 0.004 | 0.024 |
| | Within-individual variation | | 0.020 | 0.017 | 0.023 | 0.027 | 0.023 | 0.031 |

**Table S5** Variance components for the number of visits to the feeding spot in the control vs. predator treatment.

| Behavior | Variance component | Week | Control | | | Predator | | |
|---|---|---|---|---|---|---|---|---|
| | | | Estimate | Lower CI | Upper CI | Estimate | Lower CI | Upper CI |
| Visits to feeding spot | Repeatability for individual | 1 | 0.563 | 0.442 | 0.661 | 0.280 | 0.159 | 0.410 |
| | Repeatability for family | | 0.000 | 0.000 | 0.000 | 0.242 | 0.080 | 0.428 |
| | Among-individual variation in intercepts | | 2.807 | 1.772 | 4.310 | 1.015 | 0.562 | 1.635 |
| | Among-individual variation in slopes | | 0.134 | 0.084 | 0.209 | 0.186 | 0.114 | 0.296 |
| | Among-family variation in intercepts | | 0.000 | 0.000 | 0.000 | 0.878 | 0.250 | 1.968 |
| | Within-individual variation | | 2.039 | 1.737 | 2.420 | 1.526 | 1.293 | 1.828 |
| | Repeatability for individual | 2 | 0.475 | 0.364 | 0.575 | 0.275 | 0.168 | 0.407 |
| | Repeatability for family | | 0.000 | 0.000 | 0.000 | 0.223 | 0.085 | 0.408 |
| | Among-individual variation in intercepts | | 1.977 | 1.284 | 2.998 | 0.959 | 0.572 | 1.557 |
| | Among-individual variation in slopes | | 0.144 | 0.086 | 0.232 | 0.185 | 0.108 | 0.299 |
| | Among-family variation in intercepts | | 0.000 | 0.000 | 0.000 | 0.769 | 0.258 | 1.771 |
| | Within-individual variation | | 2.039 | 1.758 | 2.448 | 1.540 | 1.301 | 1.832 |
| | Repeatability for individual | 3 | 0.411 | 0.315 | 0.518 | 0.340 | 0.212 | 0.470 |
| | Repeatability for family | | 0.000 | 0.000 | 0.000 | 0.201 | 0.071 | 0.374 |
| | Among-individual variation in intercepts | | 1.560 | 1.040 | 2.318 | 1.283 | 0.795 | 2.057 |
| | Among-individual variation in slopes | | 0.159 | 0.094 | 0.269 | 0.177 | 0.108 | 0.279 |
| | Among-family variation in intercepts | | 0.000 | 0.000 | 0.000 | 0.748 | 0.227 | 1.753 |
| | Within-individual variation | | 2.042 | 1.747 | 2.448 | 1.552 | 1.321 | 1.833 |
| | Repeatability for individual | 4 | 0.404 | 0.291 | 0.512 | 0.453 | 0.321 | 0.571 |
| | Repeatability for family | | 0.000 | 0.000 | 0.000 | 0.168 | 0.061 | 0.319 |
| | Among-individual variation in intercepts | | 1.497 | 0.926 | 2.271 | 2.086 | 1.252 | 3.082 |
| | Among-individual variation in slopes | | 0.150 | 0.091 | 0.245 | 0.156 | 0.103 | 0.224 |
| | Among-family variation in intercepts | | 0.000 | 0.000 | 0.000 | 0.773 | 0.239 | 1.649 |
| | Within-individual variation | | 2.048 | 1.741 | 2.425 | 1.548 | 1.294 | 1.841 |
